# Prior scene context reshapes feature reliance during rapid perception

**DOI:** 10.64898/2026.05.10.724088

**Authors:** Sule Tasliyurt-Celebi, Benjamin de Haas, Melissa L.-H. Võ, Katharina Dobs

## Abstract

Human perception is shaped by both sensory input and prior knowledge or expectations. But how does prior contextual information influence rapid visual processing? Here, we combined eye tracking with feature-based encoding models across two experiments to predict detection latencies in a core visual task: rapid face detection in natural scenes (N = 38 per experiment). In the first experiment, we manipulated the presence of faceless scene previews. In the second experiment, we additionally restricted peripheral visual input using a moving-window paradigm, thereby increasing reliance on prior information. Across both experiments, prior context facilitated face detection, particularly for challenging images. This facilitation was already evident in the very first eye movement, suggesting that previews shape perceptual strategies from the outset. To quantify what information guided behavior, we modeled detection latencies using a set of image-based predictors capturing (i) sensory information and (ii) a scene-derived spatial prior: the expected face location. Both predictor classes explained latency variation across images. Among sensory predictors, the difference in deep neural network responses induced by the presence of the face provided the strongest out-of-sample prediction of detection latency. Critically, when scene previews were available, the contribution of the spatial prior increased, while reliance on sensory-driven features was generally reduced. Together, these findings indicate that prior scene context shifts the balance of information used for rapid face detection from sensory-driven to expectation-based spatial guidance.

## Introduction

Visual perception and cognition rely on a dynamic interplay between bottom-up and top-down signals that enable us to understand and interact with our environment. Bottom-up processes are driven by immediate sensory input, whereas top-down processes draw on prior knowledge, expectations, and experience(Firestone & Scholl, 2016; Gregory, 1970); both are shaped by task demands. This interaction has long been central to theories of visual perception, from research on visual illusions and attentional selection(Theeuwes, 2010; Treisman & Gelade, 1980) to studies of the underlying neural mechanisms(Gilbert & Li, 2013). Yet it remains unclear how the visual system combines bottom-up input with top-down signals to guide behavior(Peters et al., 2024). While some visual phenomena are characterized as fast, automatic processes driven primarily by sensory input(Alilović et al., 2019; Serre et al., 2007; Thorpe et al., 1996), growing evidence suggests that contextual expectations strongly influence visual perception, even under rapid viewing conditions(Eckstein et al., n.d.; Peelen et al., 2024; Võ & Henderson, 2010). Here, we ask how prior contextual information shapes the visual system’s reliance on sensory-driven versus expectation-based cues during a core, naturalistic visual task: the rapid detection of human faces.

This task offers a challenging test case because, in the domain of face perception, research has predominantly emphasized bottom-up processes, reflecting the fast and seemingly automatic nature of face detection(Cerf et al., 2007). Faces reliably capture attention and are the most frequently fixated objects in natural scenes(Bindemann et al., 2005; Cerf et al., 2009; de Haas et al., 2019). Humans can rapidly and accurately detect faces across a wide range of contexts(Bindemann & Lewis, 2013), with saccades to faces occurring as early as 100 ms after stimulus onset(Crouzet et al., 2010; Martin et al., 2018). Such rapid responses have often been interpreted as evidence that face detection is primarily stimulus-driven and relatively independent of top-down factors such as task demands or scene context(Crouzet & Thorpe, 2011; Prunty et al., 2024). Findings under more natural viewing conditions further support this interpretation: face-directed saccades have both lower latency and higher velocity during free viewing of complex scenes(Borovska & de Haas, 2023), and interindividual variability in rapid saccadic face responses correlates with face salience during unconstrained viewing(Broda et al., 2024). In contrast, substantial evidence indicates that face perception is neither purely stimulus-driven nor immune to top-down influences(Bindemann et al., 2010; Crouzet & Thorpe, 2011; Greene et al., 2015; Lewis & Edmonds, 2003). In the field of object perception, scene context has been shown to significantly influence object detection and perception(Aldegheri et al., 2023; Lauer & Võ, 2022). In addition, scene previews and global scene structure have been shown to guide early eye movements during visual search in natural scenes(Anderson et al., 2016; Torralba et al., 2006). Together, these findings show that contextual effects in object perception are well established, whereas evidence in face perception is mixed(Lewis & Edmonds, 2003; Nevard et al., 2023), raising the question of whether similar contextual influences also shape face detection.

To investigate the mechanisms underlying face perception, recent studies have turned to deep convolutional neural networks (CNNs) as models of human perception. CNNs have proven powerful tools for understanding human perceptual processes, including face(Dobs et al., 2022; O’Toole et al., 2018; van Dyck & Gruber, 2023) and scene(Groen et al., n.d.; Karapetian et al., 2023) perception. These models can predict primate recognition accuracy and reaction times, even for challenging images(Kar et al., 2019; Spoerer et al., 2020). Feedforward CNNs, which lack recurrent dynamics and top-down feedback, are often considered idealized models of bottom-up sensory processing(Garlichs & Blank, 2024; Peters et al., 2024), as they do not contain explicit mechanisms for integrating contextual information (even though sufficiently deep architectures may approximate some recurrent computations(Liao & Poggio, 2020)). However, training on large-scale datasets exposes these networks to statistical regularities that may implicitly encode contextual cues, posing challenges for disentangling purely sensory-driven features from learned biases, particularly in task-optimized models(Scholte & De Haan, 2025). Interestingly, even untrained CNNs have been found to contain units that respond selectively to faces(Baek et al., 2021), suggesting that architectural constraints alone may be sufficient for basic face detection, independent of visual experience. In the present study, we use both trained and untrained CNNs as computational proxies for sensory-driven features. By including untrained networks, we aim to disentangle architectural biases from effects of learned input statistics, enabling us to assess whether architectural constraints alone are sufficient to account for variance in face detection behavior.

More generally, we ask whether, and how, prior context reshapes the relative contribution of sensory-driven and expectation-based features during rapid face detection in natural scenes. To address this question, we use cross-validated encoding models across two experiments to quantify how different feature classes relate to face detection behavior, measured via eye tracking. We then assess how their contributions change with the availability of prior scene context. Sensory-driven features ranged from basic visual features (e.g., face eccentricity) and mid-level features (e.g., RMS contrast) to more abstract representational distances derived from trained and untrained CNNs. As an expectation-based feature, we included a spatial prior reflecting the expected face location within a scene. We hypothesized that prior scene previews would facilitate face detection. Specifically, we predicted that in the absence of prior context, face detection behavior would rely on sensory-driven features. In contrast, when prior contextual information was available, we expected a greater contribution of the context-based spatial prior and a reduced reliance on sensory-driven features.

## Methods

### Participants

Two independent samples of university students participated in the study. In the first experiment (face detection task), 38 participants (mean age = 26.18 years, SD = 4.05; 12 male, 26 female) with normal or corrected-to-normal vision participated. In the second experiment (moving-window task), a separate group of 38 participants (mean age = 25.00 years, SD = 3.58; 9 male, 29 female), all with normal or corrected-to-normal vision, took part. All participants provided written informed consent prior to the experiment. The sample size (N = 38 per experiment) was determined based on prior eye-tracking studies using comparable paradigms involving scene-based search tasks(David et al., 2020). In addition, an independent group of 126 participants (mean age = 23.96 years, SD = 3.48; 37 male, 89 female) participated in an online experiment (face expectation task). In this study, participants provided informed consent by clicking "Agree" at the start of the experiment. Sex/gender was determined by self-report using the response options male, female, and diverse; no participant selected the diverse option. Data on socioeconomic status (SES), communities of descent, and race/ethnicity were not collected. All experiments were conducted in accordance with the Declaration of Helsinki and approved by the local ethics committee of Justus Liebig University Giessen. Participants were compensated with either course credit or €8 per hour for their time.

#### Apparatus

Stimuli were presented on a 23.8-inch LG monitor with a resolution of 3840 × 2160 pixels and a refresh rate of 59 Hz in a dark room. Participants were seated approximately 55 cm from the monitor and remained steady throughout the experiment, using a combined forehead-and-chin rest to minimize head movement. The experiment was designed and executed using Psychtoolbox (version 3.0.16) in MATLAB R2021b(Kleiner et al., 2007) (MathWorks, Natick, MA, USA). Eye-tracking data were collected using an EyeLink 1000 system (SR Research, Ottawa, Canada).

#### Stimuli

To test the role of prior scene information on face detection behavior, we curated a dataset of 120 natural scene images, each containing a single face. These images were drawn from 12 indoor scene categories (art studio, bathroom, bedroom, bookstore, conference room, corridor, gym, kitchen, office, pharmacy, reception, and restaurant), with ten images per category. Most stimuli were sourced from the Places365 dataset(Zhou et al., 2018), while additional images were retrieved from the internet to ensure an equal number of stimuli per category. To minimize central bias and because face detection in natural scenes relies on scanning extended visual environments(Bindemann & Lewis, 2013), 82% of the stimuli featured faces located outside the image center (see Supplementary Fig. 7). These images served as target stimuli. For each target stimulus, we produced a corresponding faceless version by manually editing out the face and body using Adobe Photoshop 2022 (Adobe Inc., 2019); during this process, we also removed any associated person-related visual cues, including shadows, reflections, or other artifacts, and filled the edited regions using surrounding scene content to preserve a naturalistic appearance. These faceless images served as preview stimuli. All stimuli were displayed at a size of 46.8 × 35.1 degrees of visual angle (dva).

### Face detection task

To explore how individuals recognize faces in natural contexts and how prior scene information influences this process, participants (N = 38) performed a face detection task (Fig. 1a). Each trial began with a brief (250 ms) presentation of either a scene preview (preview condition) or a gray screen (no-preview condition). This was followed by a static noise pattern displayed for 50 ms and a fixation cross for 500 ms to minimize residual visual traces of the preview and to avoid a “face pop-out” when viewing face-present target scenes following faceless previews. The target stimulus was then presented for up to 5,000 ms, during which participants were instructed to detect the face as quickly as possible by fixating on it.

**Fig. 1.**
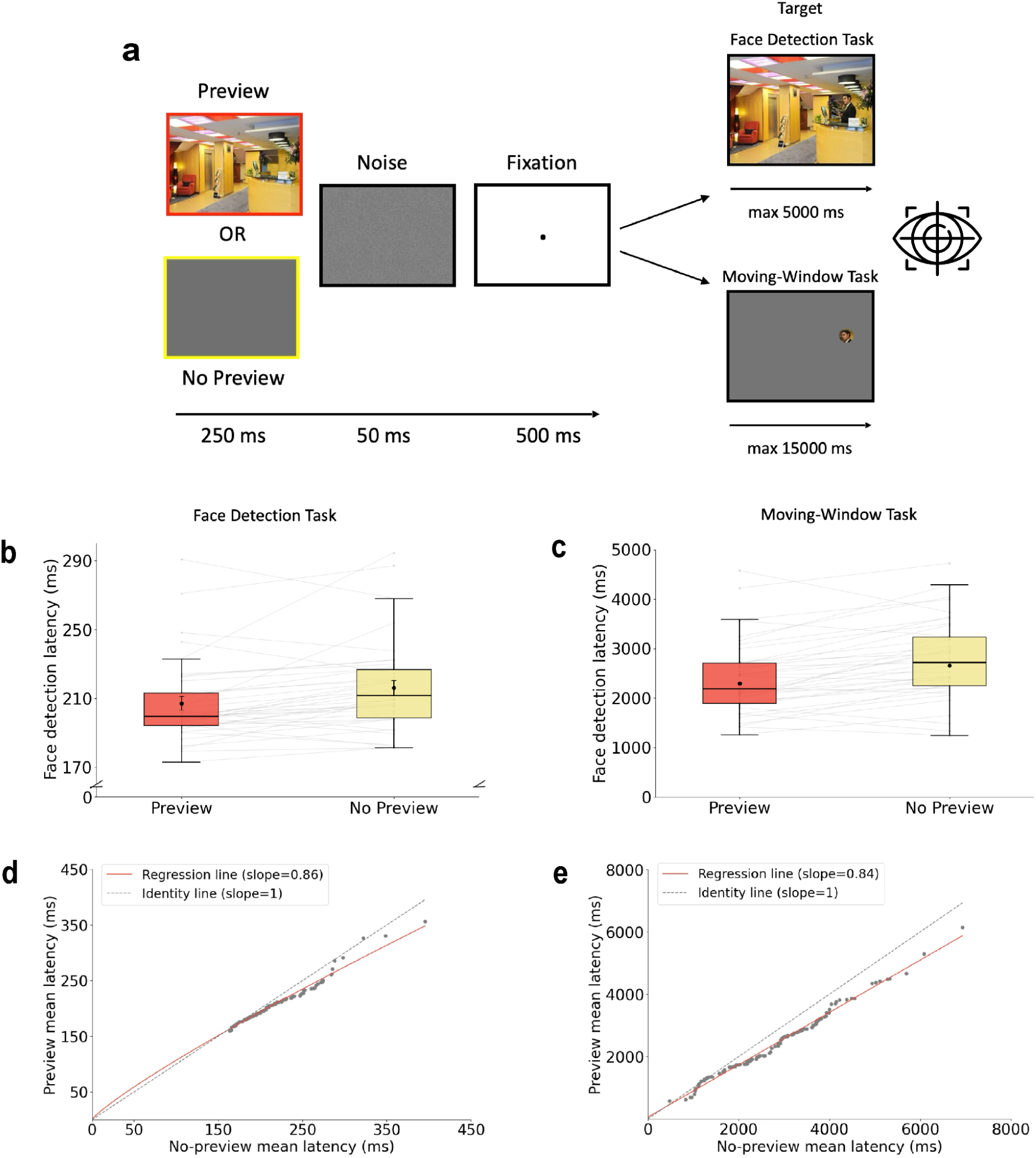
Scene previews reduce face detection latency. **a,** Task procedure and paradigms. Each trial began with either a scene preview (preview condition) or a gray screen (no-preview condition), followed by a noise mask and fixation. Participants then completed one of two face-detection paradigms: (top) a standard face detection task (N = 38), in which the full target scene was displayed (up to 5,000 ms) and participants detected the face as quickly as possible by fixating on it; or (bottom) a moving-window task (N = 38), in which only a small gaze-contingent Gaussian window revealed the scene while the remainder was masked, requiring active scanning until the first fixation on the face or a timeout (up to 15,000 ms). Face detection latency was defined as time to first fixation on the face. Images adapted from the Places365 dataset(Zhou et al., 2018), used under the MIT License. **b,** Face detection latencies were significantly shorter in the preview than in the no-preview condition (*p* < .001). **c,** in the moving-window task, face detection latencies were also significantly shorter in the preview than in the no-preview condition (*p* < .001). Gray lines represent individual participants; bars show the mean. Error bars indicate SEM. Asterisks indicate a significant difference between conditions (\*\*\**p* < 0.001; paired two-tailed t-test). **d,** Mean detection latencies in the preview condition are plotted against those in the no-preview condition for each image (face detection task). Points below the identity line (gray dashed) indicate a preview benefit. The slope of the regression line (red) was significantly less than 1 (.86; *p* < .001, permutation test), indicating that the preview advantage was larger for more difficult images. **e,** Same as in d for the moving-window task. The slope of the regression line was significantly less than 1 (.84; *p* < .001; permutation test).

Each participant completed a total of 120 trials, equally divided between the preview condition and the no-preview condition (60 trials each). The stimulus set was split into two halves, referred to as part 1 and part 2. The assignment of stimulus parts to conditions was counterbalanced across participants: half of the participants saw part 1 with preview and part 2 without preview, while the other half saw part 1 without preview and part 2 with preview. The order of stimulus presentation was randomized for each participant.

Eye-tracking data were collected to measure face detection latency, defined as the time to the first fixation on the face in the target scene. Before the experiment, a 9-point calibration and validation procedure was performed to ensure accurate gaze tracking. The experiment was divided into four blocks of 40 trials each, with three breaks between them, during which the 9-point calibration procedure was repeated. The entire experiment lasted approximately 40 minutes.

### Moving-window task

To limit the uptake of bottom-up information during search, while still manipulating and testing the effect of scene previews, participants (N = 38) performed a face detection task using a moving-window paradigm (Fig. 1a). As in the face detection task, each trial began with a 250 ms presentation of either a faceless scene preview (preview condition) or a gray screen (no-preview condition), followed by a 50 ms static noise mask and a 500 ms fixation cross. In contrast to the face detection task, visual input during target presentation was restricted to a small, gaze-contingent window rendered as a Gaussian blob (3.9° visual angle). The remainder of the display was masked with a gray screen, eliminating peripheral vision and requiring participants to actively scan the scene to locate the face. Each scene remained on screen until face detection or for a maximum of 15 seconds. Face detection latency was defined as the time to the first fixation on the face, recorded via eye tracking. All other experimental settings were identical to those in the face detection task.

### Face expectation task

We hypothesized that scene previews might enhance face detection by providing cues about where faces are likely to appear. To test this hypothesis, we conducted an additional online experiment on Pavlovia, referred to as the ‘face expectation task’. Participants (N = 126) were shown the faceless preview stimuli from the face detection task and asked to indicate the most likely location of a face within each scene by dragging a smiley-shaped mouse cursor and clicking on the predicted location (Fig. 2c). To familiarize the task, participants completed 10 practice trials before the experiment. This procedure produced face location density maps for each preview image, visualized as clusters of predicted locations (red dots in Fig. 2c). To quantify how well these predictions aligned with the actual face locations, we calculated the Euclidean distance between the mean predicted location (across participants; red dot in Fig. 2c) and the true face location in the corresponding target image (black dot in Fig. 2c). This distance served as the "face expectation" value used in subsequent analyses.

**Fig. 2.**
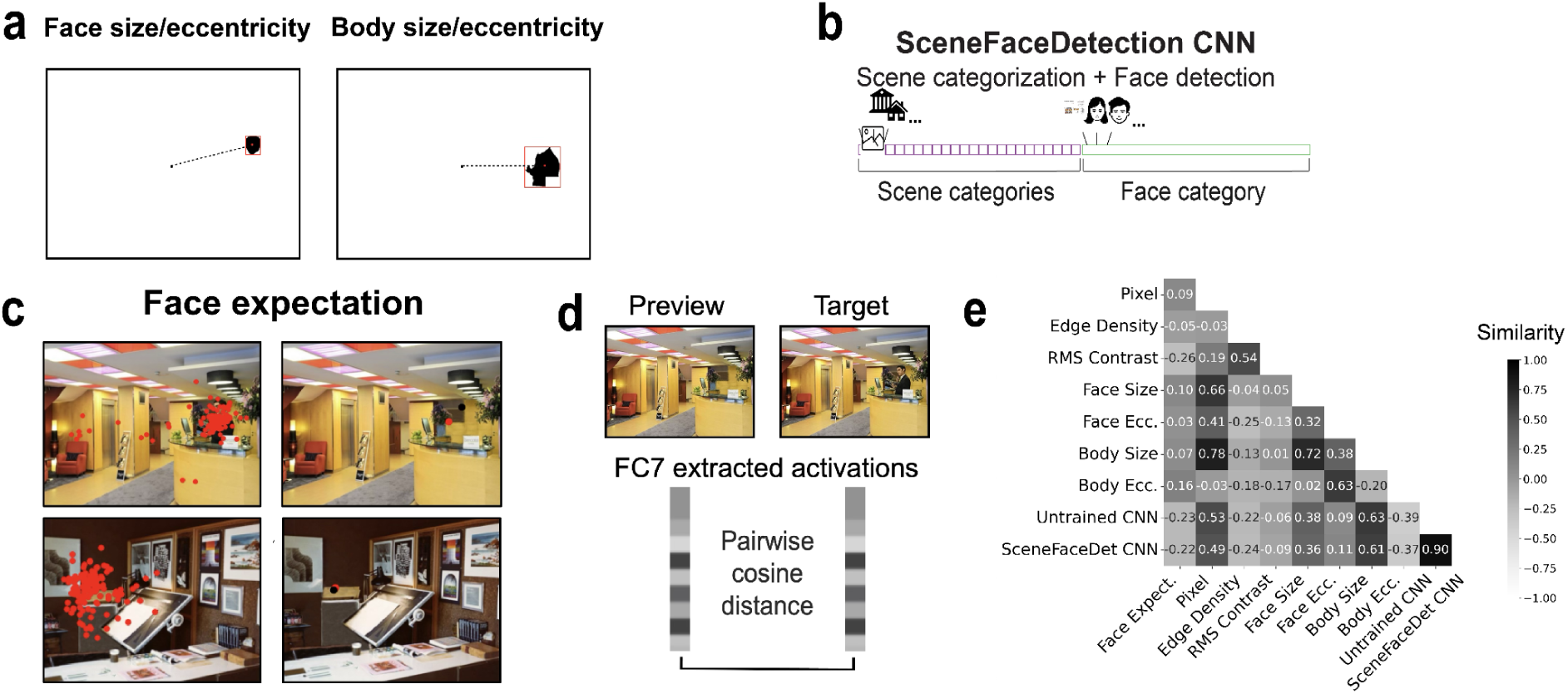
Features extraction across scenes. **a,** Example pixel masks used to compute face size and eccentricity (left) and body size and eccentricity (right). **b,** Two VGG16-based CNNs were used to derive high-level features: an Untrained CNN and a SceneFaceDet CNN trained on both scene categorization and face detection. **c,** Example density maps of expected face locations for two stimuli, visualized as clusters of predicted click locations. The Euclidean distance between the mean predicted location across participants (red dot) and the actual face location in the scene (black dot) served as the ‘face expectation’ value. Images adapted from the Places365 dataset(Zhou et al., 2018), used under the MIT License. **d,** Pairwise cosine distance between preview and target scene activations from the penultimate CNN layer (FC7) served as a feature-level change measure. **e,** Pairwise correlations (Spearman’s *r*) between all features. CNN-derived change signals were highly correlated with each other, whereas the face expectation showed low correlations with sensory-driven features.

To confirm scene categorization and perceived likelihood of face occurrence, participants also labeled each scene and provided face-likelihood ratings, though these data were not analyzed further.

### CNNs for Predicting Human Face Detection Behavior

To investigate how task-specific representations contribute to predicting human face detection behavior in natural scenes, we used two convolutional neural networks (CNNs) with VGG16 architecture(Simonyan & Zisserman, 2015) (Fig. 2b). The networks were as follows:

#### SceneFaceDet CNN

A network trained to perform both face detection and scene categorization, designed to investigate the role of face detection alongside general scene understanding in predicting human behavior.

This network categorizes 365 scene categories and a single-face category within the same classification layer, enabling concurrent processing of face and scene categorization tasks through shared representations. The SceneFaceDet CNN was trained using the Places365 dataset for 365 scene categories, each with 1160 images and the VGGFace2 dataset(Cao et al., 2018) for the single-face category. The VGGFace2 dataset included 1,423 face identities, each with 247 images. Training followed the original parameters from(Simonyan & Zisserman, 2015): stochastic gradient descent (SGD) with momentum, an initial learning rate of 10^-3^, a weight decay of 10^-4^, and momentum of 0.9. The learning rate was manually reduced to 10^-4^ and 10^-5^ when the training loss plateaued for five epochs. Cross-entropy loss was computed for random batches of 128 scene images and backpropagated to update the weights.

To improve generalization, images were scaled to a minimum side length of 256 pixels, normalized to a mean and standard deviation of 0.5, and augmented using random cropping (224 × 224 pixels), gray-scaling (20%), and horizontal flipping. To assess model performance, we evaluated the trained SceneFaceDet CNN on a held-out test set, yielding a top-5 accuracy of 0.935.

#### Untrained CNN

A randomly initialized VGG16-network used to evaluate how well features from an untrained CNN predict human face detection behavior.

#### CNN activation extraction

To evaluate each network’s ability to predict human face detection behavior, we extracted activations from the penultimate fully connected layer (FC7)—the final layer before the task-specific classification layer. These activations were computed for the 120 target scene images and their 120 corresponding faceless preview versions, mirroring the stimuli used in the human experiment. Notably, none of these images were included in the training sets of any network.

### Stimulus feature extraction and analysis

To test the role of stimulus-driven features in guiding face detection, we analyzed ten visual feature classes from our stimulus set. These included physical characteristics such as face size, body size, face eccentricity, body eccentricity, and pixel-level similarity between scenes. In order to extract features related to faces and bodies, we generated pixel masks for both face and body of each stimulus (Fig. 2a). Using these masks, we computed the size and eccentricity for both face and body. For size, we counted the total number of pixels covering the face and body. Eccentricity was defined as the distance between the centroid of the mask and the center point of the image. All calculations were performed in MATLAB R2021b(Kleiner et al., 2007) (MathWorks, Natick, MA, USA).

To capture mid-level sensory constraints related to local clutter and crowding, we extracted local RMS contrast and edge density features following established methods for quantifying clutter in natural scenes(Wallis & Bex, 2011). Images were converted to grayscale and padded to 1024 × 1024 pixels using the image mean luminance to minimize boundary artifacts. Local RMS contrast was computed as the square root of the local variance after Gaussian smoothing, and edge density was computed from a binarized Sobel edge map convolved with a Gaussian kernel. Both measures were computed at a Gaussian spatial scale of σ = 32 pixels, consistent with prior work. For each scale and feature, we sampled the feature value at the center of the face location in the padded image, yielding a scalar measure of local clutter at the target location. Log-transformed versions of these features were used in the encoding analyses. We hypothesized that the contribution of these sensory-driven features to face detection behavior would be reduced in the preview condition due to their primarily feedforward nature.

For complex visual scene- and face-specific features, we used activations from two convolutional neural networks (CNNs): the untrained CNN, and the SceneFaceDet CNN. Activations were extracted from the penultimate fully-connected layer (FC7). Additional analyses using activations from other CNN layers are provided in the Supplementary Information (Supplementary Fig. 5). For each CNN, we computed a single scalar per image pair by calculating the cosine distance between the FC7 activation vectors of the preview and target scene within the same network. This image-wise cosine distance reflects how much a given network’s representation changes between the preview and target image. That is, it captures the degree of representational change driven by the presence of a face and body in the target scene. Importantly, this measure is computed entirely within each network and does not involve any comparison of activation patterns or representational geometry across networks.

To control for low-level visual information, we conducted a pixel-level similarity analysis using the same method, substituting pixel feature vectors for layer-specific activation vectors.

As a final feature, we included the face expectation measure obtained from the face expectation experiment (Fig. 2c). For this feature, we hypothesized an enhanced contribution in the preview condition compared to the no-preview condition, reflecting the influence of prior expectations on face detection. All ten features were extracted as scalar values for each of the 120 stimuli and subsequently used as predictors in models explaining the behavioral data.

### Modeling behavioral data

#### Eye-tracking-based measurements

We used eye tracking to quantify face detection performance in both experiments through two metrics: (1) detection latency, defined as the time from target scene onset to the first fixation on the face, log-transformed prior to analysis to reduce skewness, and (2) number of fixations, defined as the number of fixations occurring from scene onset until the first fixation on the face. Additionally, to isolate presaccadic effects, we repeated the detection latency analysis on a subset of trials in which the first saccade landed directly on the face (Supplementary Note 1).

Two images were excluded from all analyses as statistical outliers, defined as exceeding three standard deviations from the condition-wise mean detection latency in either experiment. Fixations that began before stimulus onset or were shorter than 100 ms after scene onset were removed.

#### Single-feature encoding models

To examine the contribution of different feature types to face detection behavior in each experiment, we used an encoding model approach based on image-level cross-validation(Dima et al., 2022). For each feature and condition, we trained an Ordinary Least Squares (OLS) regression model to predict image-wise face detection latency from a single visual feature. The dependent variable was the mean detection latency per image, averaged across participants.

To directly test whether feature–latency relationships generalize to novel stimuli, we implemented a cross-validation procedure across images. In each fold, models were trained on a randomly selected subset of 60 images and evaluated on the remaining 60 held-out images. This procedure was repeated 100 times with different random image splits.

Model performance was quantified as the squared Spearman correlation (r²) between predicted and observed image-wise latencies in the test set. We also refer to this measure as model “predictivity” throughout this manuscript. We focus on this measure because it reflects out-of-sample generalization performance across stimuli. In contrast, regression coefficients (β) reflect the direction and scaling of feature–behavior relationships but do not capture predictive performance and are not directly comparable across features due to differences in feature scaling. Statistical significance was assessed using a permutation procedure (see Statistical analysis). To account for differences in measurement reliability between conditions, model performance was normalized by the corresponding split-half reliability estimate (see Supplementary Note 4), which provides an estimate of the noise ceiling for image-wise behavioral data by capturing the consistency of stimulus-driven responses across participants. We additionally report analyses on non-noise-corrected data (Supplementary Note 5). To isolate the unique contribution of each feature, we further performed a leave-one-feature-out analysis (Supplementary Note 6).

#### Statistical analysis

To compare eye-tracking measures (e.g., detection latency, number of fixations) between preview and no-preview conditions, we used two-tailed paired *t*-tests. Pearson correlations were used when assessing relationships within the same dataset (e.g., split-half reliability), whereas Spearman rank correlations were used for all other correlational analyses involving distinct measures.

To assess the statistical significance of encoding model performance, we used non-parametric permutation tests. Specifically, we shuffled the correspondence between observed values (i.e., detection latencies) and model predictions across stimuli and computed the correlation in the test set. This procedure was repeated 1,000 times to generate a null distribution. The two-sided *p*-value was calculated as twice the percentile rank of the observed correlation within this null distribution.

To statistically compare the predictive performance of difference features, we performed paired permutation tests on cross-validated model performance. For each comparison, we computed the difference in predictivity (ΔR²) between two features (e.g., CNN-derived predictor minus alternative feature. To generate a null distribution, we permuted the correspondence between observed and predicted latencies across stimuli (jointly for both features) and recomputed ΔR² without refitting the models. This procedure was repeated 1,000 times. Two-sided p-values were derived from the position of the observed ΔR² within the null distribution.

To assess whether prior scene context differentially modulated feature contributions, we tested for interactions using a difference-in-differences approach. Specifically, for each pair of features, we computed the difference in predictivity change across conditions (ΔR²_preview − ΔR²_no-preview) and evaluated its significance using paired permutation tests across stimuli. This allowed us to determine whether the effect of preview differed across features.

All *p*-values were Bonferroni-corrected to account for multiple comparisons.

### Sensitivity power analysis

We conducted a sensitivity power analysis in G*Power 3.1(Faul et al., 2009) for the paired-samples comparison of face detection latency between the preview and no-preview conditions in the face detection task. Assuming a two-sided *α* = .05, 80% power, and a total sample size of 38, the design was sensitive to effects of *d_z_* = .47 or larger. We report this sensitivity analysis rather than a post-hoc power analysis based on the observed effect size (Lakens, 2022). For the encoding model analyses, statistical significance was assessed using permutation testing, for which classical power analysis is not directly applicable. Model performance was evaluated across N = 120 images using 100-fold cross-validation, with a minimum detectable effect of r ≈ .18 (r² ≈ .03) based on a sensitivity analysis for a one-sample correlation (α = .05, 1 − β = .80).

### Assumption checks

Assumptions of the parametric tests were evaluated prior to analysis. For paired t-tests, the distribution of within-subject difference scores was inspected for approximate normality and the absence of extreme outliers. For OLS regression models, residuals were inspected for approximate linearity and homoscedasticity. No major violations of test assumptions were observed. Spearman correlations and permutation-based tests were used where appropriate and do not rely on the same normality assumptions as parametric tests.

### Preregistration

None of the studies reported in this manuscript were preregistered.

## Results

### Scene previews facilitate face detection performance

To test whether prior scene context affects face detection performance, we measured face detection latency via eye tracking in two experiments (Fig. 1a). In the first experiment (face detection task), participants viewed target scenes preceded either by a faceless preview of the same scene (preview condition) or by a gray screen (no-preview condition). In the second experiment (moving-window task(Castelhano & Henderson, 2007)), we restricted peripheral input using a gaze-contingent window to reduce reliance on bottom-up information.

In the face detection task, participants detected faces rapidly, with an average latency of 212 ms across conditions. Face detection latency was significantly faster in the preview than in the no-preview condition (M = 206 vs. 215 ms; *t*(37) = -3.758, *p* < .001, Cohen’s *d_z_* = -.61; 95% CI = [-.94, -.28] Fig. 1b). We did not find a significant difference in the number of fixations required to detect the face between conditions (preview: M = 1.110 ± .014 vs. no-preview: 1.129 ± .012; *t*(37) = -.526, *p* = .602, Cohen’s *d_z_*= -.09, 95% CI = [-.41, .24]; Supplementary Note 2), likely reflecting a ceiling effect, as 88% of first fixations landed directly on the face.

To assess how early this effect emerges, we repeated the latency analysis restricted to trials in which the first fixation landed on the face (Supplementary Note 1). Latencies remained significantly faster in the preview condition (*M* = 189 ms vs. 197 ms, *t*(37) = –3.394, *p* = .002; Cohen’s *d_z_* = -.55; 95% CI = [-.88, .22]), suggesting that scene previews facilitate presaccadic processing of extrafoveal content.

To test whether the observed facilitation could be explained by low-level factors, such as adaptation or face pop-out, rather than context-based guidance, we conducted a second experiment using a moving-window task that restricted the availability of bottom-up input(Castelhano & Henderson, 2007) (Fig. 1a). In this paradigm, detection latencies were again significantly faster in the preview than in the no-preview condition (M = 2,402 ms vs. 2,782 ms; *t*(37) = -5.265, *p* < .001; Cohen’s *d_z_* = -.86; 95% CI = [-1.18, -.53]; Fig. 1c), and participants required fewer fixations to detect the face (M = 6.85 vs. 8.07; *t*(37) = -6.416, *p* < .001; Cohen’s *d_z_* = -1.04; 95% CI = [-1.37, -.71]; Supplementary Note 2). We further found lower scan path ratios in the preview condition compared to the no-preview condition (*t*(37) = 3.245, *p* = .002, Cohen’s *d_z_* = -.53; 95% CI = [-.81, -.19]; Supplementary Note 3). These results suggest that, despite the increased task difficulty and restricted bottom-up input, participants consistently detected faces more quickly and efficiently when they had access to a brief scene preview. This facilitation was evident across multiple behavioral measures—including reduced latencies, fewer fixations, and more direct scan paths—indicating that participants were able to form spatial expectations that guided their search.

We next asked whether scene previews were especially beneficial for difficult images by treating no-preview latency as an index of stimulus difficulty and testing whether preview reduced latencies more for these stimuli across experiments. Plotting mean detection latencies in the preview condition against those in the no-preview condition for each stimulus revealed a strong linear relationship both in the face detection task (R² = .975, *p* < .001; Fig. 1d) and the moving-window task (R² = .990, *p* < .001; Fig. 1e), with most points falling below the identity line, suggesting overall benefit from previews. Crucially, the slope of the best-fitting line was significantly less than 1 in both the face detection task (slope = .86; *p* < .001, slope 95% CI = [.84, .88]; permutation test), and the moving-window task (slope = .84; slope 95% CI= [.82, .86]). These findings indicate that the difference between conditions increased with no-preview latency, i.e., previews conferred the largest benefit for the most difficult images.

Were these differences stable across observers? Image-wise detection latencies were highly reliable across observers in both conditions (split-half reliability; see Supplementary Note 4), supporting the analysis of stimulus-level relationships between preview and no-preview latencies. These reliability estimates were also used to normalize encoding model performance, accounting for differences in measurement noise across conditions.

### Feature contributions to face detection vary with prior context

We found that prior scene context enhanced face detection, particularly for challenging images. But how does prior context shape the extent to which different features predict detection performance? To address this question, we analyzed ten visual feature classes capturing sensory-driven and context-based information. Sensory features included basic visual properties (e.g., face size and eccentricity; Fig. 2a), mid-level measures of local clutter (RMS contrast and edge density), and representational distances derived from CNNs (Fig. 2b). For the latter, we computed representational distances between preview and target images based on penultimate layer activations (Fig. 2d), yielding a feature-level change signal. As a low-level control, we applied the same method to raw pixel values. In addition, we included a context-driven spatial prior reflecting the expected face location in each scene, derived from an independent behavioral experiment (N = 126; Fig. 2c).

Across features, correlations were generally weak to moderate (Fig. 2e), suggesting that they capture partially independent aspects of the images. CNN-derived change signals were highly correlated with each other (Spearman’s *r* > .90; *p* < .001), suggesting similar representational geometry despite differences in task optimization. Moreover, CNN-derived measures were moderately correlated with pixel-based differences (Spearman’s *r* > .49; *p* < .001) and body size (Spearman’s *r* < .61; *p* < .001), suggesting that they partially capture global image-level changes. In contrast, the face expectation measure showed weak correlations with basic visual features (Spearman’s *r* < .16; *p* = .056), and moderate negative correlations with CNN-derived measures (Spearman’s *r* < -.22; *p* = .007), indicating relative independence from sensory-driven features.

To evaluate how well each feature predicted behavior, we fit separate stimulus-level cross-validated encoding models for the preview and no-preview conditions in both experiments.

In the face detection task, multiple feature classes significantly predicted face detection latency (Fig. 3a). Among the basic visual features, all explained a significant portion of variance in both conditions: face size (preview: β = -.206 ± .057; R² = .038 ± .036; *p* < .001; no-preview: β = -.213 ± .087; R² = .066 ± .061; *p* < .001), face eccentricity (preview: β = .227 ± .081; R² = .056 ± .045; *p* < .001; no-preview: β = .378 ± .071; R² = .157 ± .064; *p* < .001), body size (preview: β = -.221 ± .077; R² = .046 ± .044; *p* < .001; no-preview: β = -.116 ± .062; R² = .106 ± .050 *p* < .001) and body eccentricity (preview: β = .400 ± .084; R² = .130 ± .055; *p* < .001; no-preview: β = .583 ± .062; R² = .342 ± .082; *p* < .001). Of these, body eccentricity was the strongest predictor overall, suggesting that body position serves as a salient spatial cue guiding gaze toward faces(Bindemann et al., 2010). To probe whether face detection latency reflects low- or mid-level image properties, we included pixel-wise similarity as a low-level control and edge density and RMS contrast as mid-level controls. Pixel-level similarity did not significantly predict face detection latency in either condition (preview: β = −.119 ± .079; R² = .023 ± .042; *p* = 1; no-preview: β = -.142 ± .082; R² = .047 ± .055; *p* = 1). Similarly, mid-level features did not significantly explain variance in face detection latency (edge density; preview: β = -.021 ± .096; R² = -.011 ± .014; *p* = 1; no-preview: β = .011 ± .088; R² = -.009 ± .021; *p* = 1; RMS contrast; preview: β = -.041 ± .084; R² = -.009 ± .016; *p =* 1; no-preview: β = -.064 ± .093; R² = .005 ± .037; *p =* 1). These results suggest that low- and mid-level image properties alone cannot account for variability in performance.

**Fig. 3.**
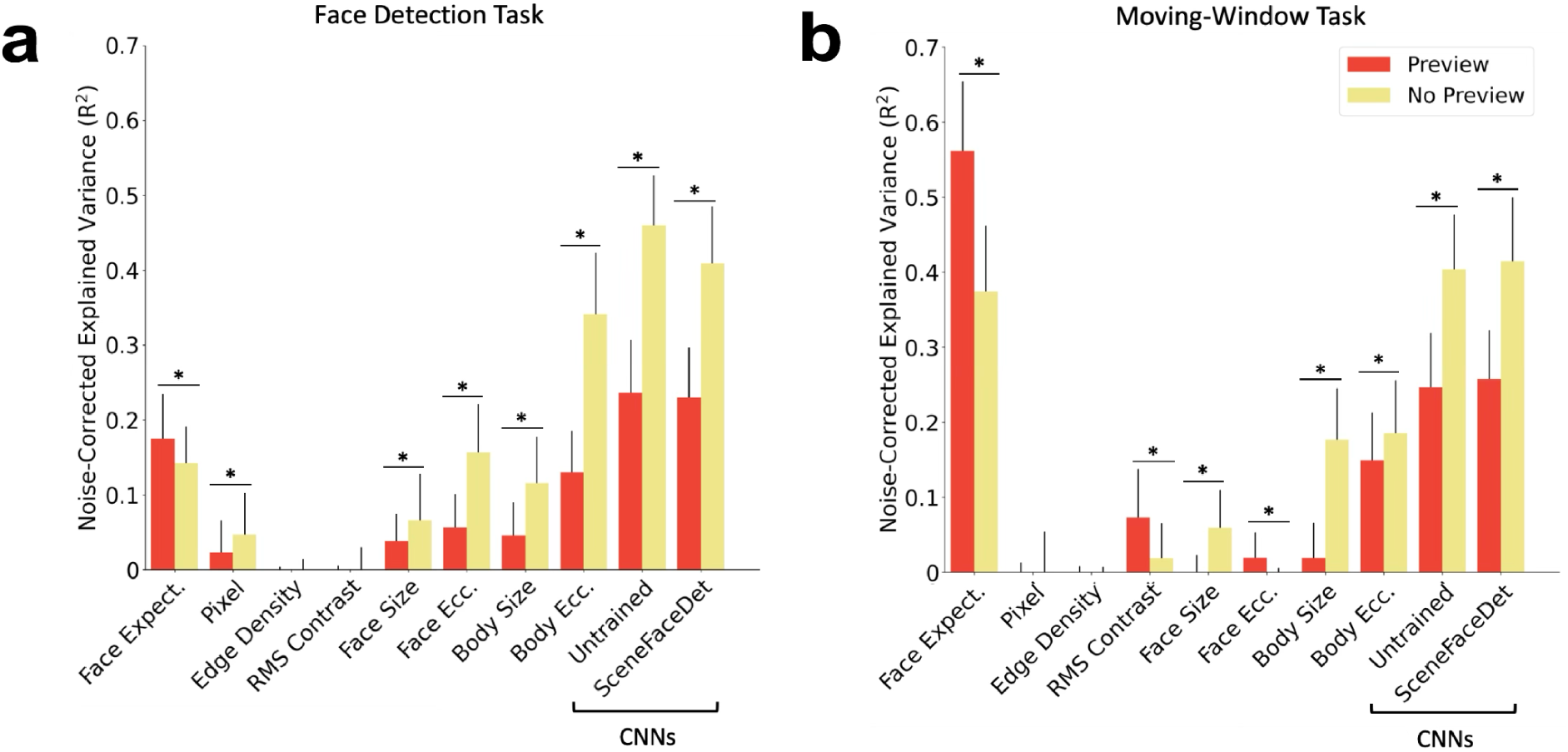
Encoding model analyses reveal feature predictivity varies with prior scene context. **a,** Noise-corrected explained variance (R²) of cross-validated single-feature encoding models predicting face detection latency from individual sensory-driven and context-driven features, shown separately for preview (red) and no-preview (yellow) conditions (N = 38). **b,** Same analysis for the moving-window task, in which visual input was restricted. The contribution of the spatial prior (face expectation) increased in the preview condition, whereas the predictivity of sensory-based features generally decreased. Error bars indicate SD across cross-validation folds. Asterisks denote significant pairwise differences (*p* < .001; permutation test).

CNN-derived change signals showed the highest overall predictivity, underscoring their relevance for modeling human visual behavior. Compared to all single-feature models, both CNN-derived predictors (SceneFaceDet CNN and Untrained CNN) explained significantly more out-of-sample variance (permutation tests on ΔR² with Bonferroni-correction; all corrected *p* < .001). Notably, the untrained CNN (preview: β = -.266 ± .056; R² = .236 ± .071; *p* < .001; no-preview: β = -.449 ± .032; R² = .460 ± .066; *p* < .001) performed comparably to the SceneFaceDet CNN (preview: β = -.297 ± .046; R² = .230 ± .066; *p* < .001; no-preview: β = -.446 ± .047; R² = .409 ± .076; *p* < .001). This result aligns with prior findings suggesting that architectural biases alone can give rise to face-selective representations(Baek et al., 2021). Predictivity increased monotonically across network depth (Supplementary Fig. 5), suggesting a hierarchical buildup of task-relevant features.

But do these CNN-derived change signals specifically reflect facial information? To test the specificity of this measure to facial information, we repeated the analysis using scenes in which the faces were occluded with gray masks (Supplementary Note 8). Explained variance dropped significantly across all networks (all *p* < 0.01), indicating that CNN-derived measures encode face-relevant information rather than generic visual changes, and that this information contributes to face detection behavior.

In addition to sensory-driven features, the context-driven spatial prior ‘face expectation’ significantly predicted detection latency in both conditions (preview: β = .338 ± .069; R² = .174 ± .060; *p* < .001; no-preview: β = .142 ± .049; R² = .125 ± .028; *p* < .001), suggesting that spatial expectations guide detection even under rapid viewing conditions.

Critically, prior scene context modulated the relative contribution of features. Across conditions, all sensory-driven features (including pixel-level, face/body, and CNN-derived change signals) showed significantly reduced predictivity in the preview condition (all *p* < .001), whereas the contribution of face expectation increased (*p* < .001). Moreover, the change in predictivity for face expectation differed significantly from that of sensory-driven features, with effects in opposite directions (all corrected *p* < .001). This pattern suggests that prior context dynamically reweights perceptual processing, reducing reliance on sensory-driven features while increasing reliance on expectation-based spatial guidance.

### Context-driven spatial guidance increases when visual input is restricted

To limit the availability of bottom-up information during search while preserving the preview manipulation, we repeated the analyses using data from the moving-window task. This yielded a similar overall pattern, with one key difference: the context-driven spatial prior (face expectation) emerged as the strongest predictor of behavior (Fig. 3b).

Basic visual features continued to significantly predict face detection latency. Among these, body eccentricity was the strongest predictor (preview: β = .335 ± .072; R² = .149 ± .063; *p* < .001; no-preview: β = .390 ± .066; R² = .185 ± .070; *p* < .001), underscoring the increased difficulty of detecting peripheral targets under restricted viewing. Body size also predicted behavior (preview: β = -.106 ± .079; R² = .019 ± .049; *p* < .001; no-preview: β = -.283 ± .084; R² = .177 ± .068; *p* < .001), although its predictivity was reduced compared to face detection task, consistent with limited access to global scene structure. Notably, face eccentricity significantly predicted behaviour only in the preview condition (β = .154 ± .074; R² = .019 ± .034; *p* < .001), but not in the no-preview condition (β = .045 ± .086; R² = -.012 ± .017; *p* = 1), suggesting that prior context may facilitate the use of spatial information that is otherwise hard to extract under constrained input. In contrast, face size significantly predicted behavior only in the no-preview condition (β = −.166 ± .087; R² = .059 ± .050, *p* < .001), but not in the preview condition (β = −.062 ± .083; R² = −.005 ± .028; *p* = 1). This complementary pattern suggests that, in the absence of prior context, observers rely more strongly on bottom-up salience cues such as size, whereas preview information enhances the use of spatial expectations.

In contrast, low- and mid-level features did not reliably predict behavior. Pixel-level changes (preview: β = -.006 ± .073; R² = -.008 ± .020; *p* < .001; no-preview: β = .-.050 ± .103; R² = -.002 ± .056; *p* = 1), and edge density (preview: β = -.002 ± .079; R² = -.013 ± .021; *p* = 1; no-preview: β = .035 ± .096; R² = -.010 ± .017; *p =* 1) showed no significant effects, while RMS contrast exhibited negative predictivity (preview: β = -.225 ± .101; R² = .073 ± .064; *p* < .001; no-preview: β = -.152 ± .093; R² = .019 ± .046; *p* < .001). These findings confirm the successful limitation of low-level image information in the moving-window task.

CNN-derived change signals remained robust predictors despite constrained visual input. Both the untrained CNN (preview: β = -.239 ± .072; R² = .246 ± .072; *p* < .001; no-preview: β = -.415 ± .074; R² = .404 ± .072; *p* < .001), and the SceneFaceDet CNN (preview: β = -.340 ± .071; R² = .257 ± .065; *p* < .001; no-preview: β = -.484 ± .076; R² = .414 ± .085; *p* < .001) explained significant variance. This suggests that CNN-derived change signals capture behaviorally relevant information even when scenes are only partially revealed, potentially reflecting differences in spatial layout or content between preview and target images.

As in the face detection task, prior scene context modulated the relative contribution of features. Across conditions, most sensory-driven features showed significantly reduced predictivity in the preview condition (all *p* < .001). In contrast, face eccentricity (*p* < .001) and RMS contrast *(p* < .001) showed small but inverted effects with their change in predictivity differing from other sensory-driven features (all corrected *p* < .001). Critically, the contribution of face expectation increased with scene previews (*p* < .001), and its change in predictivity differed from other features (all corrected *p* < .001). Moreover, the spatial prior (‘face expectation’) explained the most variance in both conditions (preview: β = .668 ± .049; R² = .562 ± .092; *p* < .001; no-preview: β = .526 ± .058; R² = .374 ± .088; *p* < .001), supporting a strong role of top-down expectations under perceptual constraints.

Taken together, these findings suggest that observers rely strongly on scene-based predictions of face location when visual input is restricted, and that prior context dynamically reweights perceptual processing, reducing reliance on sensory-driven features while increasing reliance on expectation-based spatial guidance.

## Discussion

The goal of this study was to determine whether, and how, prior scene context influences the visual system’s feature reliance during rapid visual processing. We addressed this question using two complementary experiments based on a naturalistic face detection task. Across both experiments, prior scene context, operationalized via brief previews of the faceless target scene, facilitated face detection performance, particularly for challenging images. This suggests that (i) scene previews provide meaningful cues about likely face locations, and (ii) these cues can guide early visual behavior. To examine how contextual information interacts with bottom-up input, we used feature-based encoding models to predict face detection latency from a range of visual features. In the absence of prior context, multiple sensory-driven features predicted behavior, with CNN-derived change signals showing the highest overall predictivity. However, when previews were available, the predictivity of sensory-driven features decreased, while the contribution of expectation-based cues increased. Notably, under restricted visual input in the moving-window task, the spatial prior emerged as the strongest predictor. Together, these results suggest that prior context dynamically reweights feature contributions, shifting the balance from sensory-driven to expectation-based spatial guidance during rapid face detection.

Our study provides converging evidence that top-down mechanisms shape face detection behavior, even under rapid viewing conditions. First, detection latencies were faster in the preview condition, even when the analysis was restricted to saccades that landed directly on the face. This suggests that previews facilitated presaccadic processing of extrafoveal content, providing guidance prior to the eye movement itself. Second, and more strikingly, a context-derived spatial prior, face expectation, emerged as a dominant predictor of behavior. Even though face expectation predicted face detection latency in both conditions, when prior context was available, its contribution increased significantly, while sensory-driven features were generally reduced.

These findings align with prior work showing that expected object locations can serve as effective cues during visual search(Eckstein et al., n.d.). Our results extend this work by demonstrating that spatial expectations influence visual search even under rapid viewing conditions, including tasks that are often characterized by high efficiency and speed(Crouzet & Thorpe, 2011), and that the visual system flexibly reweights its reliance on different cues depending on available context. Prior studies have shown that even brief scene previews enable rapid extraction of semantic and spatial structure(Biederman, 1981; Biederman et al., 1982; Greene & Oliva, 2009; Oliva, 2005; Oliva & Torralba, 2006), and can guide subsequent eye movements(Anderson et al., 2016; Castelhano & Henderson, 2007; Torralba et al., 2006; Võ & Henderson, 2010), particularly when preview and target share precise spatial layout. More generally, objects congruent with their surrounding context are recognized more quickly and accurately than incongruent ones(Bar, 2004; Greene et al., 2015; Krugliak et al., 2024; Lauer et al., 2020; Oliva & Torralba, 2007), in part because scene context shapes expectations not only about *what* object will appear, but also *where* it is likely to appear(Katti et al., 2017).

In our task, participants appeared to use preview-based expectations to narrow the spatial search space, resulting in faster face detection. This builds on findings that scene context predicts fixations during person detection(Ehinger et al., 2009), and extends them by quantifying how the distance between predicted and actual face locations, the spatial mismatch, modulates detection latency. This measure of spatial mismatch can be interpreted as a form of spatial prediction error, consistent with predictive coding accounts(Friston, 2010; Rao & Ballard, 1999) suggesting perception is driven by minimizing discrepancies between top-down expectations and bottom-up input. Critically, because the preview provided perfect spatial correspondence with the subsequent target scene (minus the person), the resulting spatial prior is likely scene-specific and may partly reflect narrowed spatial attention or image-specific spatial guidance. Taken together, our results show that a scene-specific spatial prior derived from scene context—the face expectation cue—is not only behaviorally relevant but emerges as the dominant feature guiding face detection when prior contextual information is available. This highlights the strength of top-down spatial expectations in shaping even rapid visual decisions, while also delineating the conditions under which such expectations are most beneficial, and reinforces the need to model visual behavior as an interplay between incoming sensory input and anticipatory contextual signals(Lotter et al., 2017).

The moving-window task further clarifies the conditions under which face expectation dominates behavior. In both paradigms, face expectation was a significant predictor in the preview and no-preview conditions, indicating that its contribution is not contingent on the presence of an explicit preview. However, face expectation was markedly stronger in the moving-window task, consistent with the constraints imposed by the gaze-contingent window. By severely limiting peripheral vision and access to global scene information, the paradigm reduces the availability and reliability of sensory-driven guidance, even when no explicit preview is provided. Importantly, the absence of a preview does not imply an absence of contextual information. Observers can still infer aspects of scene layout and canonical person locations from briefly available gist and from locally diagnostic structure encountered through the window. Under these conditions, visual search becomes more strongly guided by internally generated spatial expectations, leading face expectation to explain substantial variance even in the no-preview condition. Together, the consistency of the face expectation effect across paradigms, despite their different sensory constraints, strengthens the interpretation that this feature reflects top-down spatial guidance rather than low-level visual transients.

Our results demonstrate that all sensory-driven features—ranging from CNN-derived and pixel-level change signals to basic image-level attributes like size and eccentricity—explained less behavioral variance when prior context was available. This shift suggests that perceptual processing becomes less reliant on feedforward input when top-down information is available, even for a task as rapid and seemingly automatic as face detection. Among image-level features, body size was among the strongest predictors of behavior, and linked to CNN-derived change signals, consistent with prior findings that bodies can serve as effective cues for localizing faces(Bindemann et al., 2010; Lewis & Edmonds, 2003). Moreover, face eccentricity contributed to variation in face detection latency, indicating that faces farther from the center of the image were harder to detect—a finding in line with observers’ natural center-fixation biases(Bindemann et al., 2010). Notably, these image-level features were especially important in the absence of prior context, suggesting that when participants could not rely on scene-based expectations, they fell back on simple physical cues to guide their search(Bindemann & Burton, 2009; Lewis & Edmonds, 2003). Taken together, these findings reinforce the idea that even rapid perceptual tasks are not rigidly bottom-up. Instead, face detection reflects a flexible system that dynamically integrates available sources of information—shifting from stimulus-driven to expectation-based cues when context allows.

We find that CNN-derived change signals effectively predicted human face detection performance in natural scenes, supporting their utility as models of human visual behavior. Unlike previous studies that directly used raw CNN activations, we employed a more indirect approach: the representational distance (cosine similarity) between preview and target scenes. Despite its simplicity, this measure was among the strongest predictors of face detection latency. Notably, pixel-level similarity, used as a low-level visual control, explained behavior significantly less well. Of all the features, body size was most correlated with the CNN-derived change signal, suggesting that the CNNs may capture high-level scene properties related to body presence or scale. Moreover, when faces were masked in the stimuli, predictivity dropped, suggesting that CNN-derived signals encode face-relevant information in addition to global image differences. Together, these results highlight that human face detection behavior in natural scenes is best explained by visual representations that go beyond low- and mid-level image statistics. Accordingly, we interpret CNN-derived change signals as proxies for high-level feedforward visual representations, rather than as mechanistic models whose internal features are individually explained.

Could task-optimized CNNs exploit statistical regularities of typical face locations during training(Scholte & De Haan, 2025)? Our findings suggest otherwise: untrained CNNs—networks with randomized weights and no exposure to any visual data—predicted human behavior as well as task-optimized networks. This challenges the view that dataset-driven learning of spatial regularities is necessary to explain model performance. Instead, it suggests that architectural biases alone, even without any visual experience, can support face detection-relevant selectivity. This is consistent with recent work showing that category-selective units, including face detectors, can emerge spontaneously in untrained CNNs(Baek et al., 2021). While our face-masking analysis supports this interpretation, we cannot rule out alternative, less face-specific explanations, such as differences in global scene layout. Future work should investigate whether emergent face-selective units in untrained networks may contribute to predicting human face detection behavior. While these results underscore the power of architectural constraints, the observed reduction in their contribution under prior context also point to a limitation of purely feedforward models: their inability to incorporate dynamic expectations. Prior work has shown that explicitly modeling such expectations can enhance model performance(Katti et al., 2017). In this light, our findings highlight a promising direction for future models—combining architectural biases with flexible, context-sensitive top-down mechanisms to better account for the dynamic interplay observed in human perception(Navalpakkam & Itti, 2006).

### Limitations

While our findings shed light on how prior scene context modulates feature reliance during rapid face detection, several limitations should be acknowledged. First, our manipulation of context was based on presenting the exact same scene with and without the face, which maximized experimental control but most directly operationalizes a scene-specific spatial prior. As a result, part of the preview benefit may reflect narrowed spatial attention or image-specific spatial guidance, rather than a fully generalizable expectation formed from non-identical experiences. Importantly, this interpretation is consistent with our framing of the preview as a spatial prior that narrows the search space. Second, we used a fixed duration and timing for preview presentation, leaving open questions about how different preview durations or delays between preview and target might influence integration of prior information(Võ & Henderson, 2010). Third, our feature set was focused on hypothesis-driven features. However, incorporating data-driven or unsupervised feature representations may reveal additional structure in the data not captured by our current models(Hebart et al., 2020). Finally, while we used feedforward CNNs to model bottom-up visual processing, extending this approach to recurrent or attention-based neural network architectures could better approximate the dynamic integration of sensory and contextual information seen in the human visual system(Kar et al., 2019; Spoerer et al., 2020). Exploring these directions would not only address current limitations but also further illuminate the mechanisms by which prior knowledge shapes visual perception.

## Conclusion

Our results show that prior scene context modulates visual feature reliance during face detection in natural scenes. Even in rapid eye movement-based tasks, top-down signals—particularly expectations about likely face locations—dynamically shape visual processing. While sensory-driven features, including those captured by untrained and task-optimized CNNs, strongly predict behavior in the absence of prior information, contextual cues dominate when scene previews are available. These findings underscore the flexibility of the visual system in integrating prior knowledge to guide perception, and show how computational models can help dissociate the contributions of different feature types. By bridging behavioral, computational, and eye-tracking methods, our study advances understanding of how the brain balances sensory input and expectations in real-world vision.

## Data availability

All stimuli are available on OSF (DOI 10.17605/OSF.IO/5AYX8)(Tasliyurt-Celebi, 2025). All data will be made publicly available on GitHub (https://github.com/VCCN-lab) upon publication.

## Code availability

All Python code (version 3.8.20) used for data preprocessing, analyses, and visualization will be made publicly available on GitHub (https://github.com/VCCN-lab) upon publication.

## Author Contributions

S.T.-C., M.L.-H.V., and K.D. conceived and designed the experiments; S.T.-C. performed the experiments; S.T.-C. and K.D. analyzed the data; S.T.-C. and B.d.H. contributed materials/analysis tools; all authors wrote the paper.

## Acknowledgments

We thank Peer Herholz for his valuable contributions to reviewing, testing and updating the code. This work was supported by an ERC Starting Grant (DEEPFUNC, ERC-2023-STG-101117441), the Hessian Ministry of Science and Research, Arts and Culture (LOEWE Start Professorship), and the Deutsche Forschungsgemeinschaft (German Research Foundation, DFG) under Germany’s Excellence Strategy (EXC 3066/1 “The Adaptive Mind”, Project No. 533717223). The funding organizations had no role in the study design, data collection and analysis, decision to publish, or preparation of the manuscript.

