## Supplementary material for "Prior scene context reshapes feature reliance during rapid perception": SI

This file includes:

Supplementary Notes

Supplementary Methods

### Supplementary Note 1

#### Face detection latency with first fixations landing on the face

To examine whether the benefit of scene previews persists even when participants detect the face with a single fixation, we analyzed face detection latency restricted to the subset of trials in which the first fixation landed directly on the face (88% of trials). Even within this subset, participants detected faces significantly faster in the preview condition compared to the no-preview condition (*M* = 189 ms vs. 197 ms, $t(37) = –3.39$, $p = .002$, Cohen's $d_z = -.55$, 95% CI = [-.88, -.22]). This result suggests that scene previews help presaccadic processing of extrafoveal content, facilitating detection even before the eyes land on the face.

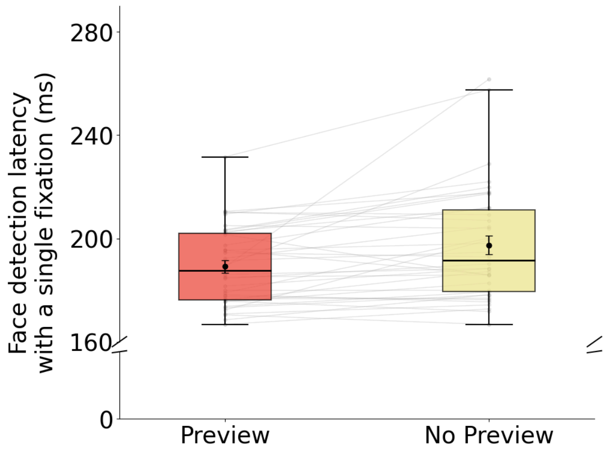


**Supplementary Fig. 1 |** Face detection latencies were significantly faster in the preview than in the no-preview condition (M = 189 ms vs. 197 ms; $p = .002$, Cohen's $d_z = -.55$, 95% CI = [-.88, -.22]), even when restricting the analysis to only first fixations that landed directly on the face. Gray lines show individual participant data. Error bars indicate SEM.

### Supplementary Note 2

#### Number of fixations

To assess whether the preview benefit reflects changes in search efficiency (i.e., how many gaze shifts were needed to locate the face), we compared the number of fixations required to detect the face across conditions. In the standard face detection task, detection typically occurred immediately after image onset, and the number of fixations did not reliably differ between preview and no-preview trials (preview: $M$ = 1.110 ± 0.014; no-preview: $M$ = 1.129 ± 0.012; $t(37) = -.526$, $p = .602$, Cohen's $d_z$ = -.09, 95% CI = [-.41, .24]), consistent with the idea that previews primarily modulate how quickly the face is detected rather than the number of discrete fixations needed. In contrast, under gaze-contingent viewing, where peripheral information is strongly reduced and participants must actively scan, previews substantially reduced the number of fixations required to find the face (preview: $M$ = 6.95 vs. no-preview: $M$ = 8.21; $t(37) = 6.416$, $p < .001$; Cohen's $d_z$ = -1.04; 95% CI = [-1.37, -.71]). This pattern suggests that scene previews improve search guidance when exploration is necessary, reducing the number of fixations required to detect the face.

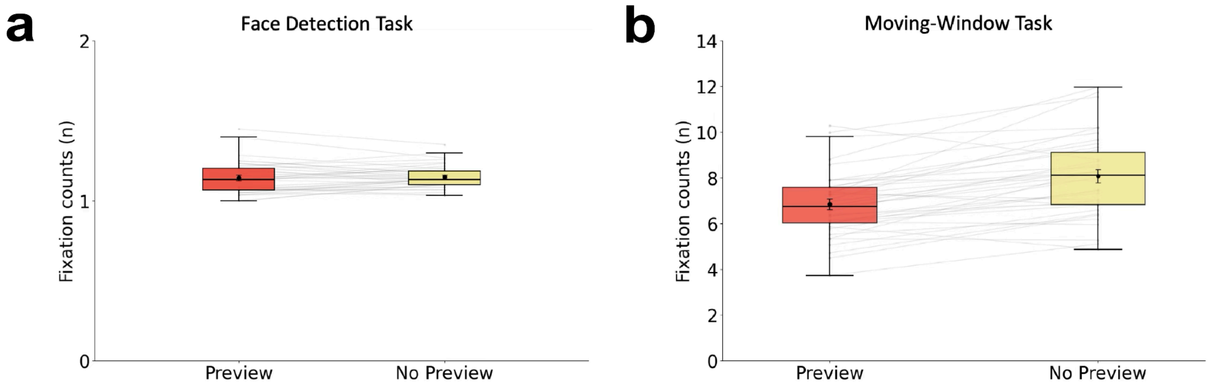


**Supplementary Fig. 2 | Number of fixations. a,** The number of fixations required to detect the face did not differ significantly between conditions (preview: M = 1.110 ± 0.014 vs. no-preview: M = 1.129 ± 0.012; $t(37) = -.526$, $p = .602$, Cohen's $d_z$ = -.09, 95% CI = [-.41, .24]). **b,** Participants required fewer fixations to detect the face in the preview condition (M = 6.95 vs. 8.21; $p < .001$, Cohen's $d_z$ = -1.04; 95% CI = [-1.37, -.71]). Gray lines show individual participant data. Error bars indicate SEM.

### Supplementary Note 3

#### Scan path ratio

To further assess how prior scene context modulates search efficiency, we computed the scan path ratio for each trial in the moving-window task. This analysis was restricted to the moving-window task as the face detection task did not involve a sufficient number of saccades to compute meaningful scan paths. This metric quantifies the efficiency of eye movement paths by comparing the total distance covered by the eyes (i.e., the sum of all saccade lengths from scene onset to the first fixation on the face) to the shortest possible straight-line distance from the central fixation point to the center of the face. A scan path ratio of 1 reflects a perfectly direct route to the target, whereas larger values indicate more circuitous search behavior. Consistent with previous definitions[1,2], this measure captures the strength of search guidance prior to target fixation, independent of post-fixation decision processes.

Scan path ratios were significantly lower in the preview condition compared to the no-preview condition ($t(37) = 3.245$, $p = .002$, Cohen's $d_z = -.53$; 95% CI = [-.81, -.19]), indicating more direct and efficient search when participants had access to a scene preview. This result complements the latency and fixation count findings, reinforcing the notion that scene previews facilitate more efficient, goal-directed visual search under constrained viewing.

### Supplementary Note 4

#### Internal consistency

To assess the reliability of image-wise detection latencies, we computed split-half consistency separately for the preview and no-preview conditions. For each condition, participants were randomly split into two groups 100 times. Within each split, mean detection latencies were computed across participants for each image in both halves, and Pearson correlations were calculated between the resulting vectors. The resulting correlation values were then corrected using the Spearman-Brown formula to yield final reliability estimates. Consistency was high in both conditions (preview: $r = .94$; no-preview: $r = .90$; Spearman-Brown corrected), suggesting that image-specific detection patterns are highly reliable and consistent across observers. We repeated the same split-half procedure for the moving-window task. Consistency was likewise high in both conditions (preview: $r = .79$; no-preview: $r = .83$; Spearman–Brown corrected), indicating that image-wise latency patterns remain reliable under gaze-contingent viewing.

In addition, we computed a single shared noise ceiling for each experiment. The estimated split-half reliability, measured as the Spearman–Brown-corrected split-half reliability of image-wise detection latencies, was $r = .96$ for the face detection task and $r = .89$ for the moving-window task.

### Supplementary Note 5

**Single-feature encoding models reveal context-dependent predictivity (uncorrected)**

As a complement to the noise-corrected analyses reported in the main text, we repeated the same single-feature encoding analyses using uncorrected explained variance (Supplementary Fig. 5). The overall pattern closely matched the main results.

In the face detection task, basic visual features significantly predicted detection latency in both conditions, including face size (preview: β = -.206 ± .057; $R^2$ = .036 ± .034; $p$ < .001; no-preview: β = -.213 ± .087; $R^2$ = .060 ± .055; $p$ < .001), face eccentricity (preview: β = .227 ± .081; $R^2$ = .053 ± .042; $p$ < .001; no-preview: β = .378 ± .071; $R^2$ = .142 ± .058; $p$ < .001), body size (preview: β = -.221 ± .077; $R^2$ = .043 ± .041; $p$ < .001; no-preview: β = -.116 ± .062; $R^2$ = .105 ± .056; $p$ < .001), and body eccentricity (preview: β = .400 ± .084; $R^2$ = .122 ± .052; $p$ < .001; no-preview: β = .583 ± .062; $R^2$ = .309 ± .074; $p$ < .001). Pixel-level similarity did not significantly predict face detection latency in either condition (preview: β = -.119 ± .079; $R^2$ = .006 ± .010; $p$ = 1; no-preview: β = -.142 ± .082; $R^2$ = .010 ± .014; $p$ = 1). Similarly, edge density (preview: β = -.018 ± .096; $R^2$ = -.009 ± .013; $p$ = 1; no-preview: β = .032 ± .088; $R^2$ = -.007 ± .019; $p$ = 1) and RMS contrast (preview: β = -.038 ± .084; $R^2$ = -.009 ± .015; $p$ = 1; no-preview: β = -.076 ± .095; $R^2$ = .006 ± .033; $p$ = 1) did not significantly predict behavior. By contrast, both CNN-derived features explained comparatively high variance, including the untrained CNN (preview: β = -.266 ± .056; $R^2$ = .221 ± .068; $p$ < .001; no-preview: β = -.449 ± .032; $R^2$ = .416 ± .060; $p$ < .001) and the SceneFaceDet CNN (preview: β = -.297 ± .046; $R^2$ = .215 ± .062; $p$ < .001; no-preview: β = -.446 ± .047; $R^2$ = .369 ± .069; $p$ < .001). Face expectation also significantly predicted detection latency in both conditions (preview: β = .338 ± .069; $R^2$ = .163 ± .056; $p$ < .001; no-preview: β = .142 ± .049; $R^2$ = .128 ± .045; $p$ < .001).

A similar pattern was observed in the moving-window task. Face eccentricity significantly predicted behavior only in the preview condition (β = .154 ± .074; $R^2$ = .015 ± .027; $p$ < .001), but not in the no-preview condition (β = .045 ± .086; $R^2$ = -.010 ± .04; $p$ = 1). Face size significantly predicted behavior only in the no-preview condition (β = -.166 ± .087; $R^2$ = .049 ± .042; $p$ < .001), but not in the preview condition (β = -.062 ± .083; $R^2$ = -.004 ± .022; $p$ = 1). Body size (preview: β = -.106 ± .079; $R^2$ = .015 ± .037; $p$ < .001; no-preview: β = -.283 ± .084; $R^2$ = .146 ± .057; $p$ < .001) and body eccentricity (preview: β = .335 ± .072; $R^2$ = .119 ± .050; $p$ < .001; no-preview: β = .390 ± .066; $R^2$ = .154 ± .058; $p$ < .001) significantly predicted performance in both conditions. Pixel-level similarity did not significantly predict behavior in both conditions (preview: β = -.006 ± .073; $R^2$ = -.006 ± .016; $p$ = 1; no-preview: β = -.050 ± .103; $R^2$ = -.001 ± .046; $p$ = 1). Edge density again did not significantly predict behavior (preview: β = -.002 ± .079; $R^2$ = -.010 ± .017; $p$ = 1; no-preview: β = .035 ± .096; $R^2$ = -.008 ± .014; $p$ = 1), whereas RMS contrast showed a significant effect (preview: β = -.225 ± .101; $R^2$ = .058 ± .051; $p$ < .001; no-preview: β = -.152 ± .093; $R^2$ = .015 ± .038; $p$ < .001). Both CNN-based predictors again explained substantial variance, including the untrained CNN (preview: β

= -.239 ± .072; $R^2$ = .196 ± .057; $p$ < .001; no-preview: β = -.415 ± .074; $R^2$ = .335 ± .060; $p$ < .001) and the SceneFaceDet CNN (preview: β = -.340 ± .071; $R^2$ = .204 ± .052; p < .001; no-preview: β = -.484 ± .076; $R^2$ = .348 ± .071; $p$ < .001). Face expectation was the strongest predictor in both conditions (preview: β = .668 ± .049; $R^2$ = .446 ± .073; $p$ < .001; no-preview: β = .526 ± .058; $R^2$ = .310 ± .073; $p$ < .001).

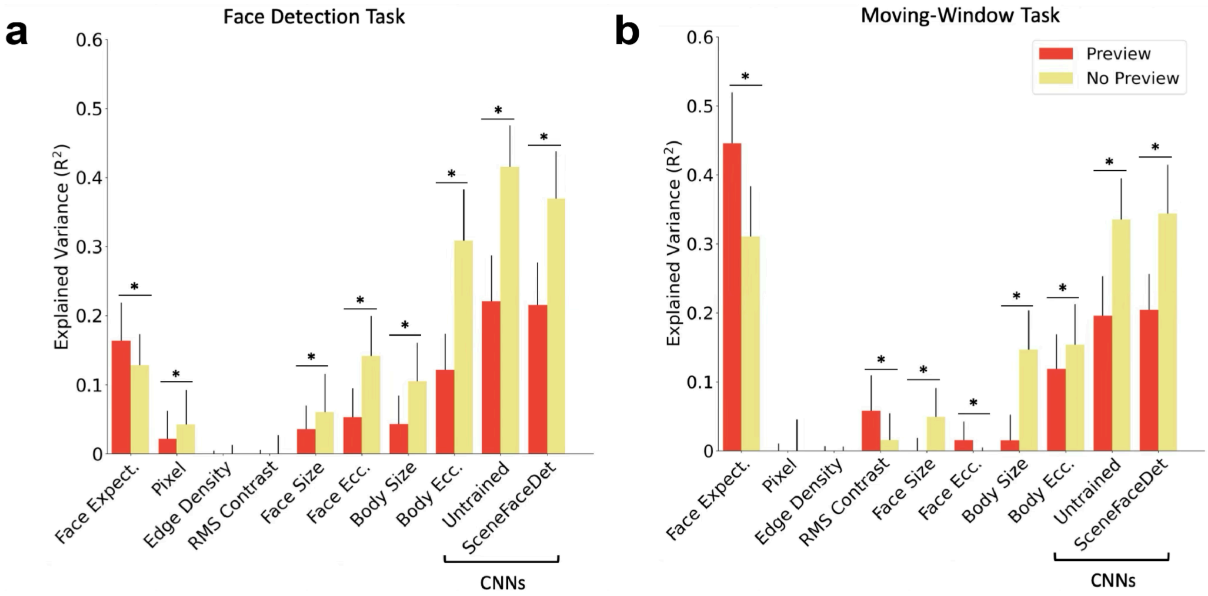


**Supplementary Fig. 3 | Encoding model analyses reveal feature predictivity varies with prior scene context (uncorrected). a,** Explained variance ($R^2$) of cross-validated single-feature encoding models predicting human face detection latency (both N = 38 participants) from individual sensory-driven and context-driven features, shown separately for preview (red) and no-preview (yellow) conditions, without noise correction. **b,** Same analysis for the moving-window task. Error bars indicate SDs across cross-validation folds. Asterisks indicate significant pairwise differences ($p$ < .001; permutation test).

### Supplementary Note 6

**Unique feature contributions support the role of face expectation**

To isolate the unique contribution of each feature, we performed a leave-one-feature-out analysis: we subtracted the variance explained by a reduced model (excluding the feature of interest) from that of the full model (including all features). Prior to model fitting, all predictors were standardized to ensure comparability across features and to avoid scaling-related biases in variance estimates. This approach estimates the unique variance accounted for by each feature beyond what is captured by all others. The resulting pattern diverged notably from the single-feature encoding models.

In the no-preview condition, several features explained unique variance in detection latency. Among the basic visual features, face eccentricity was the only feature that significantly accounted for unique variance ($R^2$ = .038 ± .037; $p$ < .001), while face size ($R^2$ = -.001 ± .033; $p$ = 1), body size ($R^2$ = -.002 ± .023; $p$ = 1) and body eccentricity ($R^2$ = -.020 ± .029; $p$ = 1) did not contribute uniquely. This suggests that the position of the face, rather than its size, plays a key role in driving detection latency. Using a single CNN-derived predictor, the SceneFaceDet CNN accounted for significant unique variance, with a stronger unique contribution in the no-preview than the preview condition ($R^2$ = .058 ± .029; $p$ < .001). Pixel-level similarity did not contribute uniquely to detection latency ($R^2$ = .005 ± .023, $p$ = .49). Likewise, the mid-level sensory features edge density ($R^2$ = -.004 ± .020; $p$ = 1) and RMS contrast did not contribute uniquely to detection latency ($R^2$ = -.017 ± .24; $p$ = 1; Supplementary Fig. 6a). Interestingly, the context-based feature face expectation explained the largest amount of unique variance ($R^2$ = .042 ± .044, $p$ < .001), suggesting that even without explicit scene previews, observers form spatial expectations that influence face detection latency[42,10].

In contrast, in the preview condition, the unique contribution of most features was markedly reduced. Face expectation was the strongest unique predictor ($R^2$ = .090 ± .044, $p$ < .001), and the SceneFaceDet CNN also significantly contributed unique variance, although its effect was considerably smaller ($R^2$ = .009 ± .016; $p$ < .001). This pattern suggests that expectation-based cues play a dominant role when prior context is available, with residual contribution from high-level CNN-derived representations All other features did not explain unique variance (face size; $R^2$ = .004 ± .035; $p$ = 1; face eccentricity; $R^2$ = -.003 ± .017; $p$ = 1; body size; $R^2$ = -.016 ± .026; $p$ = 1; body eccentricity; $R^2$ = -.010 ± .026; $p$ = 1; pixel-level similarity; $R^2$ = -.008 ± .016; $p$ = 1; edge density; $R^2$ = -.006 ± .016; $p$ = 1; RMS contrast; $R^2$ = -.010 ± .021; $p$ = 1). This result further supports the conclusion that prior scene context dynamically reweights the contributions of visual features, reducing reliance on input-driven cues and increasing reliance on expectation-based, top-down information.

To test whether the same pattern holds under constrained visual input, we applied the same unique-variance analysis to the moving-window task (Supplementary Fig. 6b). As in the face detection task, face expectation uniquely accounted for the largest variance across both conditions (preview: $R^2 = .275 \pm .073$; $p < .001$; no-preview: $R^2 = .171 \pm .059$; $p < .001$). In contrast, basic visual features contributed relatively little unique variance, particularly in the preview condition. Specifically, face eccentricity ($R^2 = .011 \pm .037$; $p = .043$) and body eccentricity ($R^2 = .026 \pm .039$; $p < .001$) contributed significant variance in the no-preview condition but not in the preview condition (face eccentricity; $R^2 = -.015 \pm .031$; $p = 1$; body eccentricity; $R^2 = -.015 \pm .028$; $p = 1$). Face size (preview; $R^2 = -.003 \pm .026$; $p = 1$; no-preview; $R^2 = .001 \pm .025$; $p = X$),) and body size (preview; $R^2 = -.014 \pm .024$; $p = 1$; no-preview; $R^2 = -.021 \pm .034$; $p = 1$) showed no unique contribution in either condition.

Using a single CNN-derived predictor, the SceneFaceDet CNN showed a significant unique contribution in both conditions (preview: $R^2 = .017 \pm .037$; $p < .001$; no-preview: $R^2 = .051 \pm .049$; $p < .001$). However, this effect was modest relative to the context-based predictor, suggesting that CNN-based representations account for only limited unique variance under constrained visual input.

Neither pixel-level similarity explained only small unique variance in the no-preview condition (preview: $R^2 = -.008 \pm .033$; $p = 1$; no-preview: $R^2 = .032 \pm .044$; $p < .001$), nor edge density (preview: $R^2 = -.013 \pm .027$; $p = 1$; no-preview: $R^2 = -.017 \pm .035$; $p = .1$) or RMS contrast (preview: $R^2 = -.007 \pm .015$; $p = 1$; no-preview: $R^2 = -.009 \pm .019$; $p = .1$) accounted for unique variance in detection latency.

In sum, the moving-window paradigm provided a stringent test of face detection under highly constrained visual conditions. Encoding analyses revealed that while some sensory-driven features still contributed to performance, the strongest and most unique predictor of detection behavior was a spatial prior derived from scene context: face expectation. These results suggest that prior scene context shapes perceptual guidance, even in rapid perceptual tasks. More broadly, they highlight the utility of contextual information in shaping visual search behavior when perceptual input is limited.

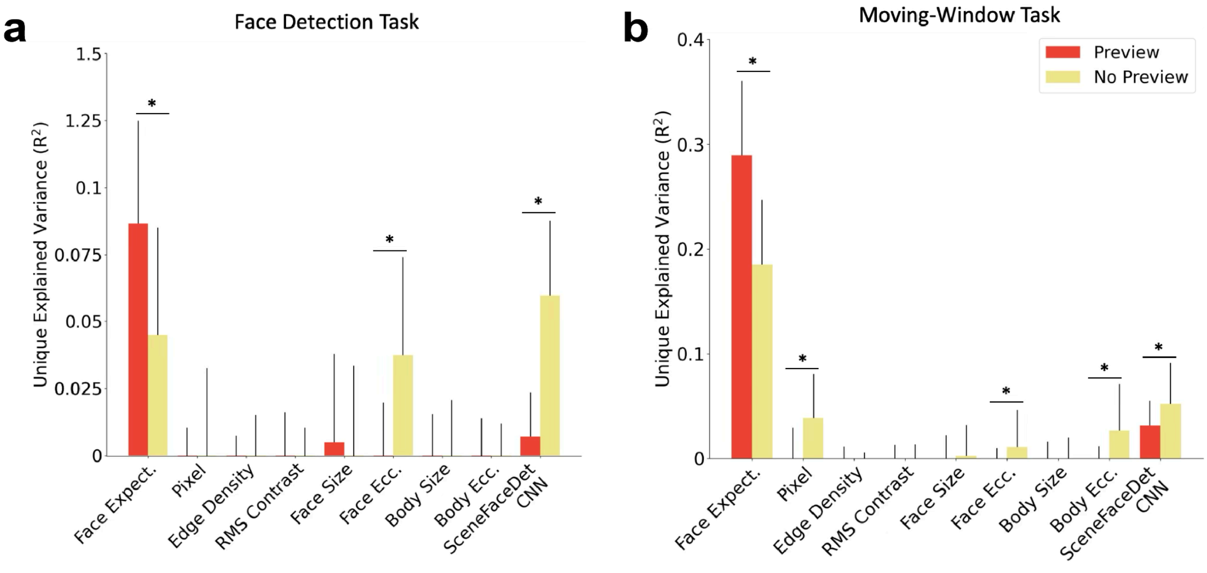

**Supplementary Fig. 4 | Unique Explained Variance.** Unique explained variance, quantified as the reduction in $R^2$ when each feature is removed from the full multivariate encoding model, indicating variance in detection latency uniquely attributable to that feature beyond all others. **a.** Face detection task **b.** Moving-window task (both N = 38 participants). Error bars indicate SDs across cross-validation folds. Asterisks indicate significant pairwise differences ($p < .01$; permutation test).

### Supplementary Note 7

#### Layerwise CNN predictivity analysis

To assess whether the predictive power of the CNNs increases with layer depth, we computed the Spearman rank correlation between explained variance and layer depth for each model, separately for the preview and no-preview conditions (Supplementary Fig. 5a-b). We observed a significant positive correlation in all cases: Preview condition: Untrained CNN: r = .857; $p < .014$ and SceneFaceDet CNN: r = 1; $p < .001$; No-preview condition: Untrained CNN: r = .893; $p < .007$ and SceneFaceDet CNN: r = .964; $p < .001$).

These results indicate that deeper layers consistently show greater predictivity, supporting the idea that high-level representations—rather than low-level features—are most relevant for modeling face detection behavior.

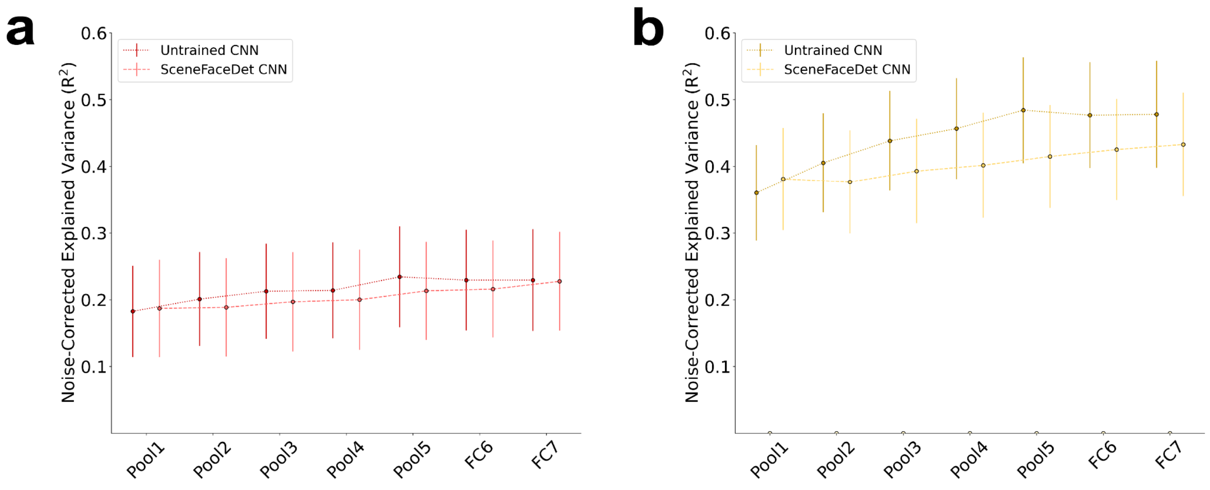


**Supplementary Fig. 5 | Layer-wise CNN predictivity. a,** Explained variance across all layers of the CNNs in the preview condition. ***b*,** Explained variance across all layers of the CNNs in the no-preview condition. Error bars indicate SD across cross-validation splits.

### Supplementary Note 8

#### Testing the contribution of facial information to CNN predictivity

To assess the contribution of facial information to CNN-based predictions, we occluded the face region in each target stimulus using a gray mask (Supplementary Fig. 8a). We then computed pairwise cosine distances between the FC7 activations of the preview and the face-masked target scenes.

High-level features derived from CNNs showed the highest overall predictivity for face detection (Supplementary Fig. 4b-c). When we repeated the analysis using masked versions of the target scenes, explained variance dropped significantly for all networks in both conditions (preview: untrained CNN: face visible: $\beta = -.266 \pm .056$; $R^2 = .221 \pm .068$; $p < .001$; face masked: $\beta = -.120 \pm .088$; $R^2 = .078 \pm .101$; $p < .001$ and SceneFaceDet CNN: face visible: $\beta = -.297 \pm .046$; $R^2 = .215 \pm .062$; $p < .001$; face masked: $\beta = -.040 \pm .121$; $R^2 = .008 \pm .122$; $p = .22$; for no-preview: untrained CNN: face visible: $\beta = -.449 \pm .032$; $R^2 = .416 \pm .060$; $p < .001$; face masked: $\beta = -.262 \pm .084$; $R^2 = .239 \pm .010$; $p < .001$ and SceneFaceDet CNN: face visible: $\beta = -.446 \pm .047$; $R^2 = .369 \pm .069$; $p < .001$; face masked: $\beta = -.126 \pm .113$; $R^2 = .122 \pm .182$; $p < .001$).

All comparisons were statistically significant ($p < .001$; permutation test). These results confirm that the CNN-derived features reflect sensitivity to facial information, and that masking the face substantially reduces the networks' ability to predict human detection behavior.

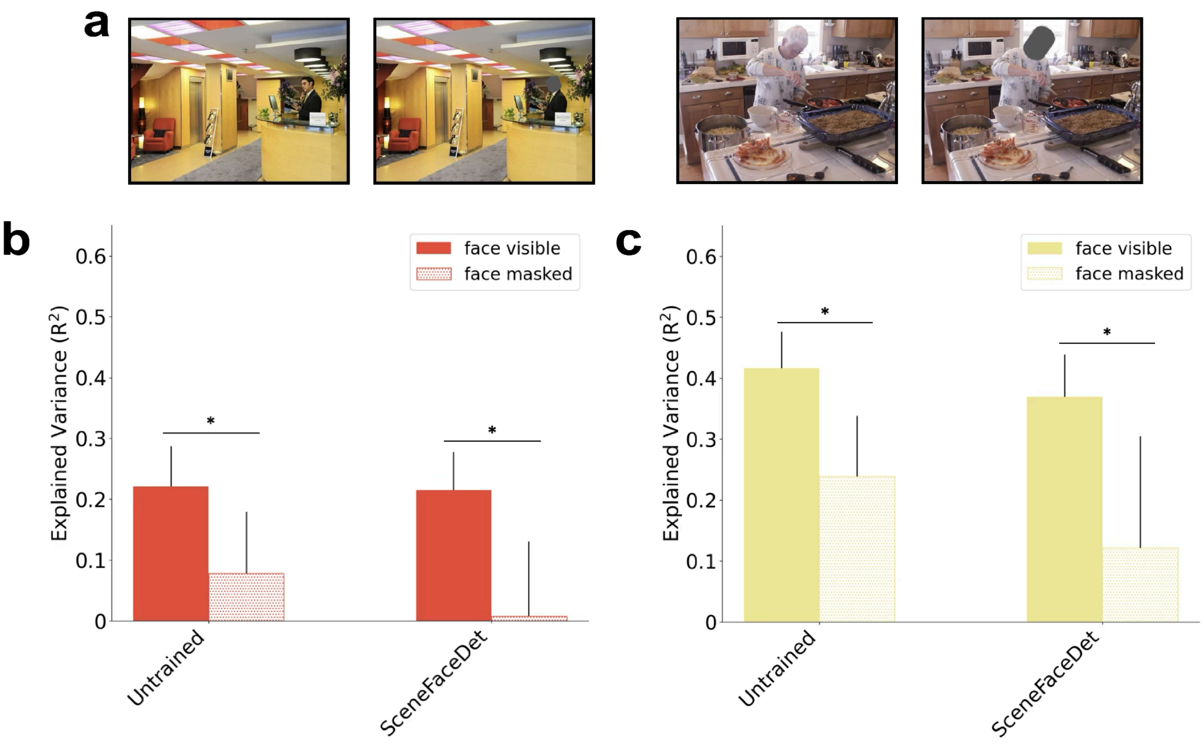


**Supplementary Fig. 6 | Face occlusion analysis. a,** Example stimulus pairs showing the original target image (left) and the face-masked version (right), where the face was occluded with a gray circle. **b,** Explained variance for each model in the preview condition, comparing target scenes with visible faces (solid bars) versus masked faces (dotted bars). **c,** Same comparison for the no-preview condition. Asterisks indicate significant pairwise differences ($p$ < .01; permutation test). Error bars represent SD across cross-validation splits.

### Supplementary methods

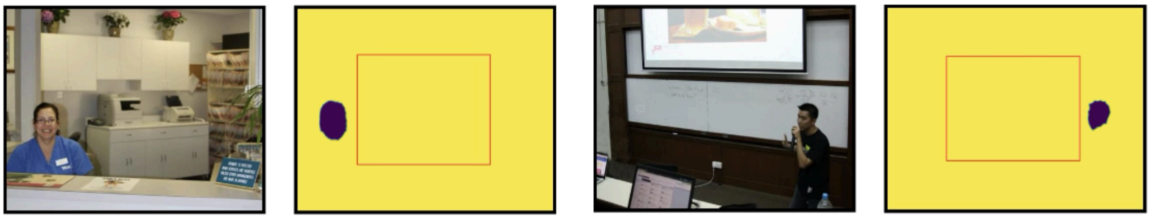

**Supplementary Fig. 7 | Example face locations.** In 82% of the stimuli, faces were located outside the center of the image (indicated by the red rectangle).

**Supplementary Fig. 1 |** Face detection latencies were significantly faster in the preview than in the no-preview condition (M = 189 ms vs. 197 ms; $p$ = .002, Cohen's $d_z$ = -.55, 95% CI = [-.88, -.22]), even when restricting the analysis to only first fixations that landed directly on the face. Gray lines show individual participant data. Error bars indicate SEM.

**Supplementary Fig. 2 | Number of fixations. a,** The number of fixations required to detect the face did not differ significantly between conditions (preview: M = 1.110 ± 0.014 vs. no-preview: M = 1.129 ± 0.012; $t$(37) = -.526, $p$ = .602, Cohen's $d_z$ = -.09, 95% CI = [-.41, .24]). **b,** Participants required fewer fixations to detect the face in the preview condition (M = 6.95 vs. 8.21; $p$ < .001, Cohen's $d_z$ = -1.04; 95% CI = [-1.37, -.71]). Gray lines show individual participant data. Error bars indicate SEM.

**Supplementary Fig. 3 | Encoding model analyses reveal feature predictivity varies with prior scene context (uncorrected). a,** Explained variance ($R^2$) of cross-validated single-feature encoding models predicting human face detection latency (both N = 38 participants) from individual sensory-driven and context-driven features, shown separately for preview (red) and no-preview (yellow) conditions, without noise correction. **b,** Same analysis for the moving-window task. Error bars indicate SDs across cross-validation folds. Asterisks indicate significant pairwise differences ($p$ < .001; permutation test).

**Supplementary Fig. 4 | Unique Explained Variance.** Unique explained variance, quantified as the reduction in $R^2$ when each feature is removed from the full multivariate encoding model, indicating variance in detection latency uniquely attributable to that feature beyond all others. **a.** Face detection task **b.** Moving-window task (both N = 38 participants). Error bars indicate SDs across cross-validation folds. Asterisks indicate significant pairwise differences ($p$ < .01; permutation test).

**Supplementary Fig. 5 | Layer-wise CNN predictivity. a,** Explained variance across all layers of the CNNs in the preview condition. ***b*,** Explained variance across all layers of the CNNs in the no-preview condition. Error bars indicate SD across cross-validation splits.

**Supplementary Fig. 6 | Face occlusion analysis. a,** Example stimulus pairs showing the original target image (left) and the face-masked version (right), where the face was occluded with a gray circle. **b,** Explained variance for each model in the preview condition, comparing target scenes with visible faces (solid bars) versus masked faces (dotted bars). **c,** Same comparison for the no-preview condition. Asterisks indicate significant pairwise differences ($p < .01$; permutation test). Error bars represent SD across cross-validation splits.

**Supplementary Fig. 7 | Example face locations.** In 82% of the stimuli, faces were located outside the center of the image (indicated by the red rectangle).
